# Mass Action Kinetic Model of Apoptosis by TRAIL-Functionalized Leukocytes

**DOI:** 10.1101/300400

**Authors:** Emily E. Lederman, Michael R. King

## Abstract

**Background:** Metastasis through the bloodstream contributes to poor prognosis in many types of cancer. A unique approach to target and kill colon, prostate, and other epithelial-type cancer cells in the blood has been recently developed that causes circulating leukocytes to present the cancer-specific, liposome-bound Tumor Necrosis Factor (TNF)-related apoptosis inducing ligand (*TRAIL*) on their surface along with *E* – *selectin* adhesion receptors. This approach, demonstrated both in vitro with human blood and in mice, mimics the cytotoxic activity of natural killer cells. The resulting liposomal TRAIL-coated leukocytes hold promise as an effective means to neutralize circulating tumor cells that enter the bloodstream with the potential to form new metastases.

**Results:** The computational biology study reported here examines the mechanism of this effective signal delivery, by considering the kinetics of the coupled reaction cascade, from TRAIL binding death receptor to eventual apoptosis. In this study, a collision of bound TRAIL with circulating tumor cells (CTCs) is considered and compared to a prolonged exposure of CTCs to soluble TRAIL. An existing computational model of soluble TRAIL treatment was modified to represent the kinetics from a diffusion-limited 3D reference frame into a 2D collision frame with advection and adhesion to mimic the *E* – *selectin* and membrane bound TRAIL treatment. Thus, the current model recreates the new approach of targeting cancer cells within the blood. The model was found to faithfully reproduce representative observations from experiments of liposomal TRAIL treatment under shear. The model predicts apoptosis of CTCs within 2 hr when treated with membrane bound TRAIL, while apoptosis in CTCs treated with soluble TRAIL proceeds much more slowly over the course of 10 hrs, consistent with previous experiments. Given the clearance rate of soluble TRAIL in vivo, this model predicts that the soluble TRAIL method would be rendered ineffective, as found in previous experiments.

**Conclusion:** This study therefore indicates that the kinetics of the coupled reaction cascade of liposomal *E* – *selectin* and membrane bound TRAIL colliding with CTCs can explain why this new approach to target and kill cancer cells in blood is much more effective than its soluble counterpart.

## 2 Background

Cancer metastasis accounts for more than 90% of cancer-related deaths (1). In many types of cancer, circulating tumor cells are shed from the primary tumor site into peripheral circulation where they then can extravasate into extravascular space to form metastatic tumors. (2–4). Recent studies have shown that CTCs from primary tumors express sialyated carbohydrate ligands which interact with selectins on the surface of the endothelium (5, 6). These selectins can begin to tether to the sialyated carbohydrate ligands, in a fashion similar to leukocyte interaction with endothelium. These rapid force-dependent binding interactions can trigger rolling adhesion and eventually firm adhesion to the endothelium, facilitating survival and formation of micrometastases (7–9). Surgery and radiation, while proven effective in treating primary tumors, pose challenges due to the limited detectability of distant micrometastases.

New methods to target CTCs have been developed in vivo and hold promise in reducing the metastatic load and the formation of new tumors. One recent technology uses leukocytes as a drug delivery mechanism. Leukocytes and CTCs are similar in size and rigidity, causing both to migrate to the near wall region of blood vessels. For every CTC, there are ∼1 × 10^6^ leukocytes circulating, which effectively surround the CTC, making leukocytes an attractive carrier for cancer drug delivery (10–13).

It has been shown that functionalizing leukocytes with liposomes decorated with *TRAIL* (*memTRAIL*) and *E* – *selectin* (*ES*), an adhesion molecule, is an effective way of treating circulating cancer cells in flowing human blood in vitro, and in the peripheral circulation of mice in vivo (12, 14). This method of treatment is more effective than soluble *TRAIL* (*sTRAIL*); however, the mechanism of this enhanced apoptosis response has not yet been fully elucidated. Beyond concentrating *memTRAIL* in the close vicinity of CTCs, there are two other key reasons why this method of treatment is believed to be so effective. First, the shearing caused by blood can help to promote the collision of *memTRAIL* with CTC, effectively increasing the on-rate of binding. Previous studies have shown that increased shear has a direct correlation with the sensitivity of cancer cells to *TRAIL* (15). Secondly, it is possible that *E* – *selectin* briefly tethers the liposome to the CTC after collision, effectively reducing the slip velocity after collision and lowering the off-rate of *TRAIL* binding *DRs.*

Several models have been built to gain a better quantitative understanding of the reaction cascade pathway that takes place when tumor cells exposed to *sTRAlL* undergo apoptosis, but previous models have not considered the *memTRAIL* interacting with CTCs in a tethered 2-D frame of reference. Some important considerations which must be captured in such a model are 2-D binding reaction kinetics, the effects of a slip velocity, and the effects of cell adhesion. This, in turn, will better represent the case of leukocytes functioned with *memTRAIL*, in a shearing blood flow, with *E* – *selectin* temporaril tethering CTCs to the treated leukocytes. Our model builds off and significantly extends Albeck et al.’s model which captures the coupled reaction pathway by numerically integrating reaction rate laws via MATLAB’s ordinary differential equation (ODE) solver (16). During apoptosis, the potent effector caspase 3 (*C*3) is activated by extracellular stimuli such as *TRAIL. C*3 degrades the proteome and activates DNAses, which dismantle chromosomes of cells committed to die (17). Caspase activation represents an irreversible change in cell fate regulated by the assembly of complexes on death receptors, binding of pro- and anti-apoptotic members of the *Bcl* – 2 family to each other in cytosolic and mitochondrial compartments, mitochondria-to-cytosol translocation of *Smac* and cytochrome c (*CyC*), and the direct repression of caspases by inhibitor apoptosis proteins (IAPs)(18–24). In the ODE-based model of *C*3 regulation, the mass action kinetics of a typical CTC undergoing apoptosis are captured to better understand this *memTRAIL* model by examining the interplay of each reagent’s concentration within the reaction cascade as a function of time. From this, new insights are revealed to explain why the sheared *memTRAIL* model is notably more effective in inducing apoptosis in CTCs.

## 3 Results

### 3.1 Specific Reactant Concentration Profiles

Specific values of *TRAIL*, *DR*, *k*_+_ and *k*_ were used for the following four cases of *TRAIL* binding to *DRs: sTRAIL*, *memTRAIL* without shear, *memTRAIL* with shear but without adhesion, and *memTRAIL* with both shear and adhesion. These four cases were chosen to mimic experiments carried out by Mitchell et. al, which showed that *TRAIL* was most potent when TRAIL and *E* – *selectin* were tethered to the surface of a liposome, and sheared during treatment of tumor cells(14). It has been suggested that *E* – *selectin* helps promote the binding of *TRAIL* and *DR* by causing the liposomes to adhere to CTCs(14). This effect of adhesion on *memTRAIL* binding *DR* was included in the model.

#### 3.1.1 *cPARP* instantaneous concentrations

Each concentration profile was normalized with its maximum concentration, to better examine the relative time progression of the reaction pathway rather than the relative concentrations within the reaction pathway. When focusing on cleavage of *PARP*, *memTRAIL* under shear with adhesion induced apoptosis the fastest of all of the treatment methods, at *T_d_* = 1.8 *hr* (Figure 1). This value is very close to that found experimentally by Mitchel et al, where there was a 98% reduction in reported circulating tumor cells, after allowing the *ES/TRAIL* treatment to circulate in mice for 2.5 hrs, when compared with mice treated with *ES* alone (14). Following that, the second fastest treatment method was *memTRAIL* without adhesion in shear at *T_d_* = 3.5 *hr. memTRAIL* without shear and *sTRAIL* had much longer times to apoptosis at *T_d_* = 12.5 *hr* and *T_d_* = 10.5 *hr* respectively.

**Figure 1.**
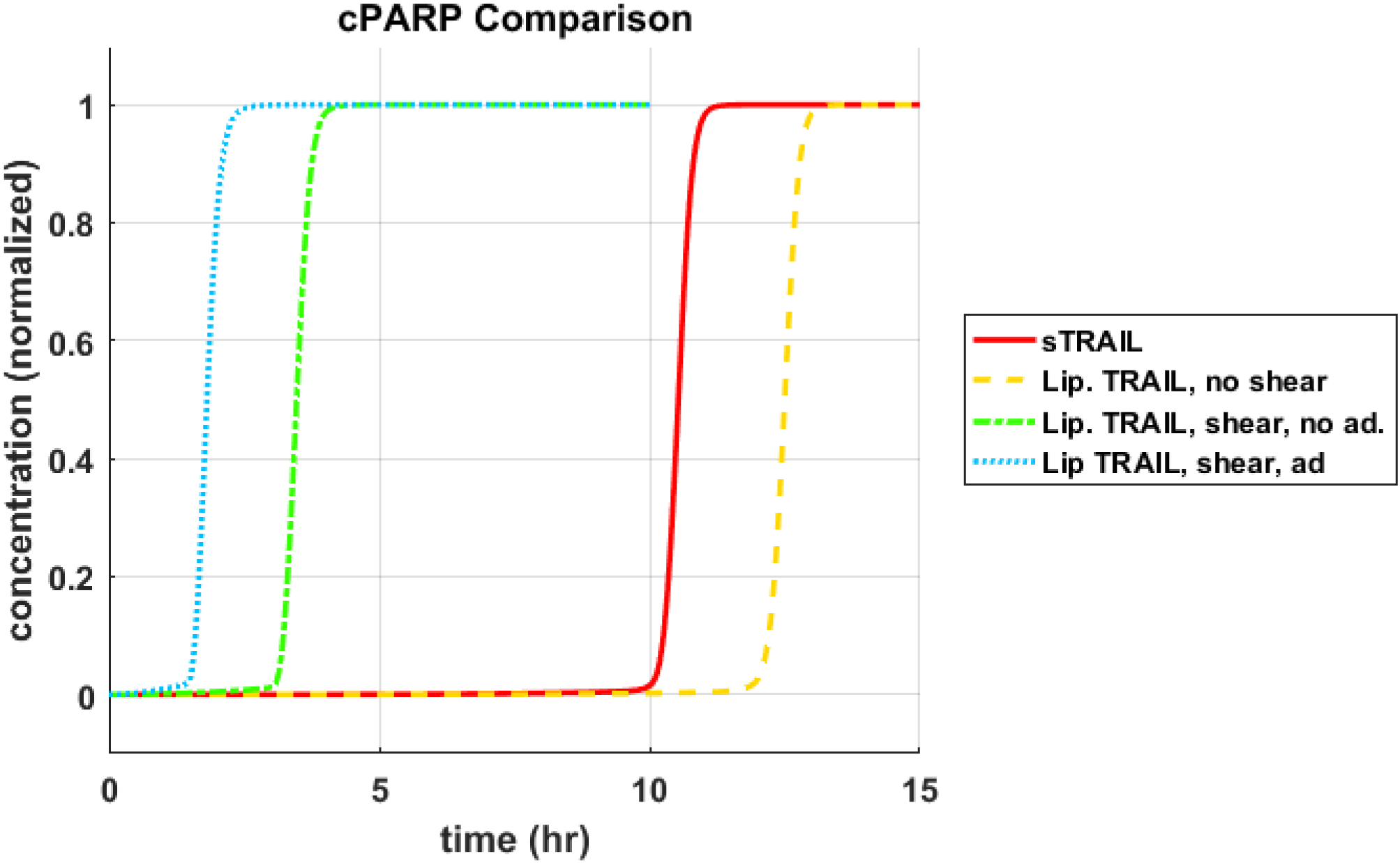
Time until apoptosis T_d_ as a function of (a) TRAIL and DR concentration, (b) TRAIL concentration and k_+_ and (c) TRAIL cocncentration and k_.

To reveal the underlying mechanism at work here, the pathway was divided into two sections, pre-mitochondrial pathway and post-mitochondrial pathway. Within these two sections, the essential reactants’ concentration were plotted as a function of time to identify why sheared *memTRAIL* delivery is a more efficient method of drug delivery both in simulations and experimentally.

#### 3.1.2 Pre-mitochondrial Species Concentration Profiles

It was observed that the pre-mitochondrial pathway transition (marked by a sudden graded response in the reactant concentrations) coincided with the cleavage of *PARP* for higher values of *T_d_* (Figure 2 a-c); however, this happened before the transition of *cPARP* for smaller values of *T_d_* (Figure 2d). It was also observed that an initial change in reagent concentration was present for the *memTRAIL* case with shear and adhesion (Figure 2d) but less prominent for the other treatment methods. These observations suggest that the pre-mitochondrial pathway is activated much faster for *memTRAIL* with adhesion and shear than for the other treatment methods.

**Figure 2.**
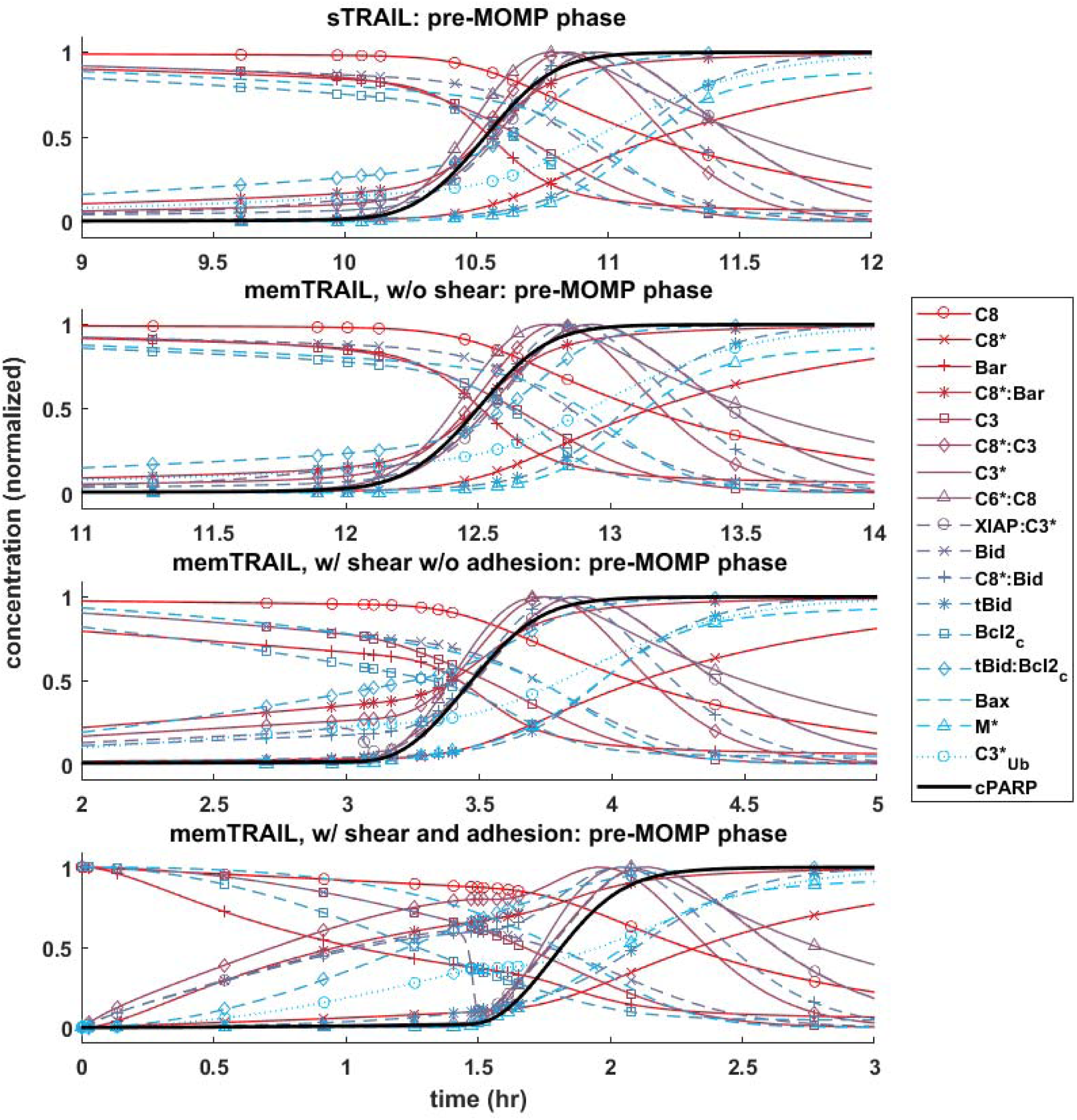
Concentration profile ofcPARP as a function of time for 4 different cases of TRAIL binding to DRs.

#### 3.1.3 Post-mitochondrial Species Concentration Profiles

Next, the post mitochondrial pathway, which transitions during and after the permeabilization of the mitochondrial outer membrane, was considered. It was observed that the reagents underwent a sharper transition as *T_d_* decreased for different treatment methods, indicating that this pathway is less inhibited by upstream reagents for *memTRAIL* delivery with shear and adhesion (Figure 3).

**Figure 3.**
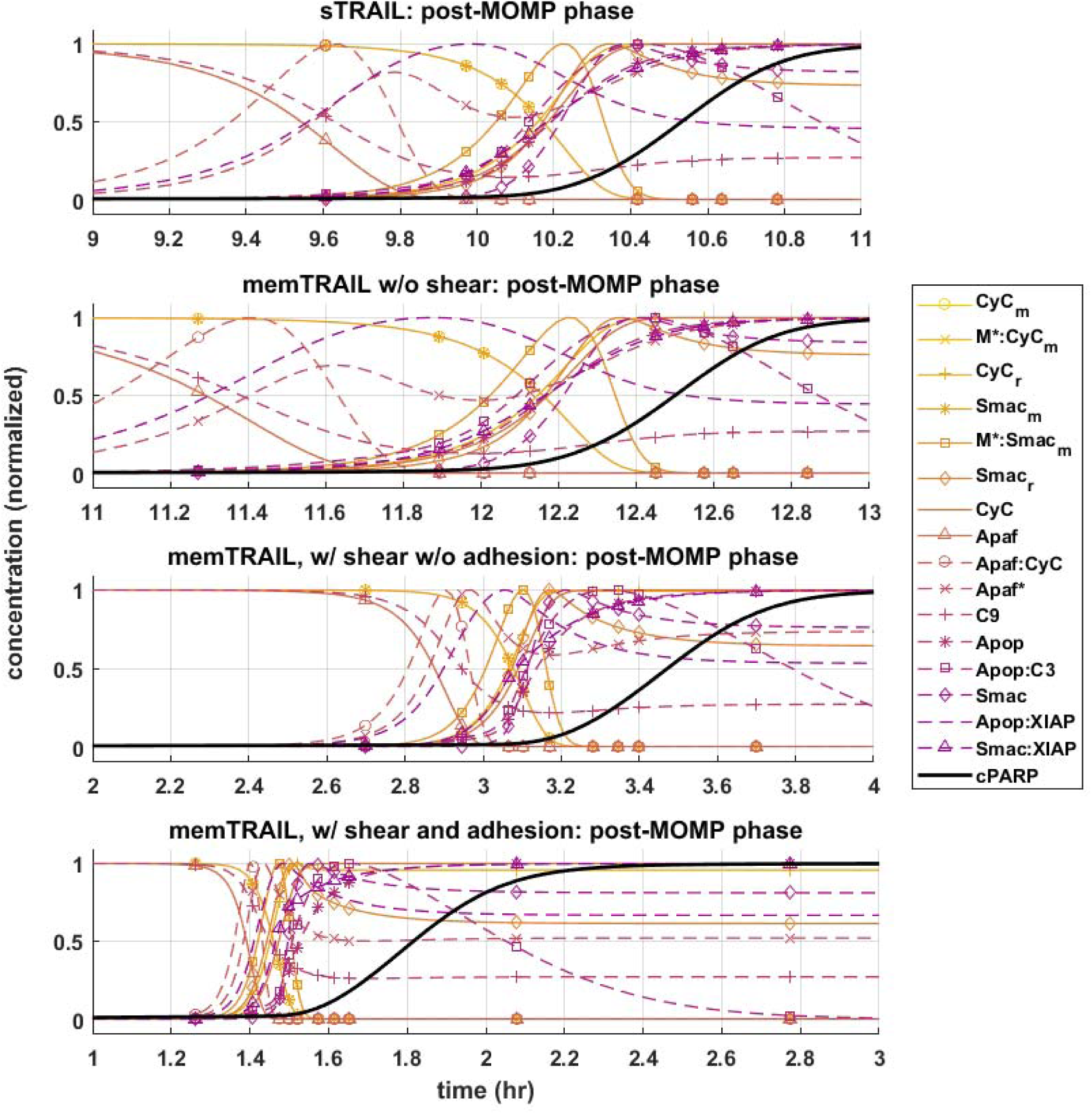
Pre-mitochondrial pathway species concentration profiles as a function of time for four different cases of TRAIL binding to DRs: sTRAIL, memTRAIL without shear, memTRAIL with shear and without adhesion, and memTRAIL with shear and adhesion.

#### 3.1.4 C3 and XIAP Concentration Profiles

Given these observations regarding the pre and post mitochondrial membrane reaction pathways, we next considered where the two pathways meet. This sheds light on the specific mechanism that allows for faster apoptosis in CTCs exposed to *memTRAIL* with shear and adhesion. Two species are of greatest importance in this analysis: *C*3 and *XIAP.* To simplify the comparison, only two critical cases were considered – *sTRAIL* and *memTRAIL* with shear and cell adhesion. Given that the time until apoptosis had already been quantified and that we were most interested in how each reactant profile emerges with respect to *cPARP’s* transition, *T_d_* was subtracted from the time vector to re-center each image around the time of cell death. The curves were normalized to the maximum concentration of reagent for *sTRAIL* pathway. In cases where the reagent concentration is relatively higher for reactants within the *memTRAIL* pathway, then the profile displays values greater than 1, and if relatively less, then values less than 1.

It is evident that a sudden increase emerges in concentration of species *C*8^∗^: *C*3, *XIAP*: *C*3^⋆^, and
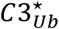
for the liposomal TRAIL method (Figure 4b). One notable difference in the relative quantities of reagents. *C*8^∗^: *C*3, *Apop*: *C*3, *C*3^∗^, *XIAP*: *C*3^∗^, and *Apop*:*XIAP* comparing the two cases is that all had lower maximum concentrations than their *sTRAIL* pathway counterparts, while *C*3^∗^: *PARP* and *XIAP*: *C*3^∗^ peaked at higher values. Another notable difference between the two conditions is the order in which reagents emerge. *Apop*:*XIAP* and *XIAP* transitioned before other reagents for the *sTRAIL* pathway, while *C*8^∗^: *C*3, *XIAP*: *C*3^∗^ and
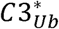
had already completed 40-75% of their respective transitions by the time *Apop*:*XIAP* and *XIAP* began to transition for the *memTRAIL* pathway (Figure 4).

**Figure 4.**
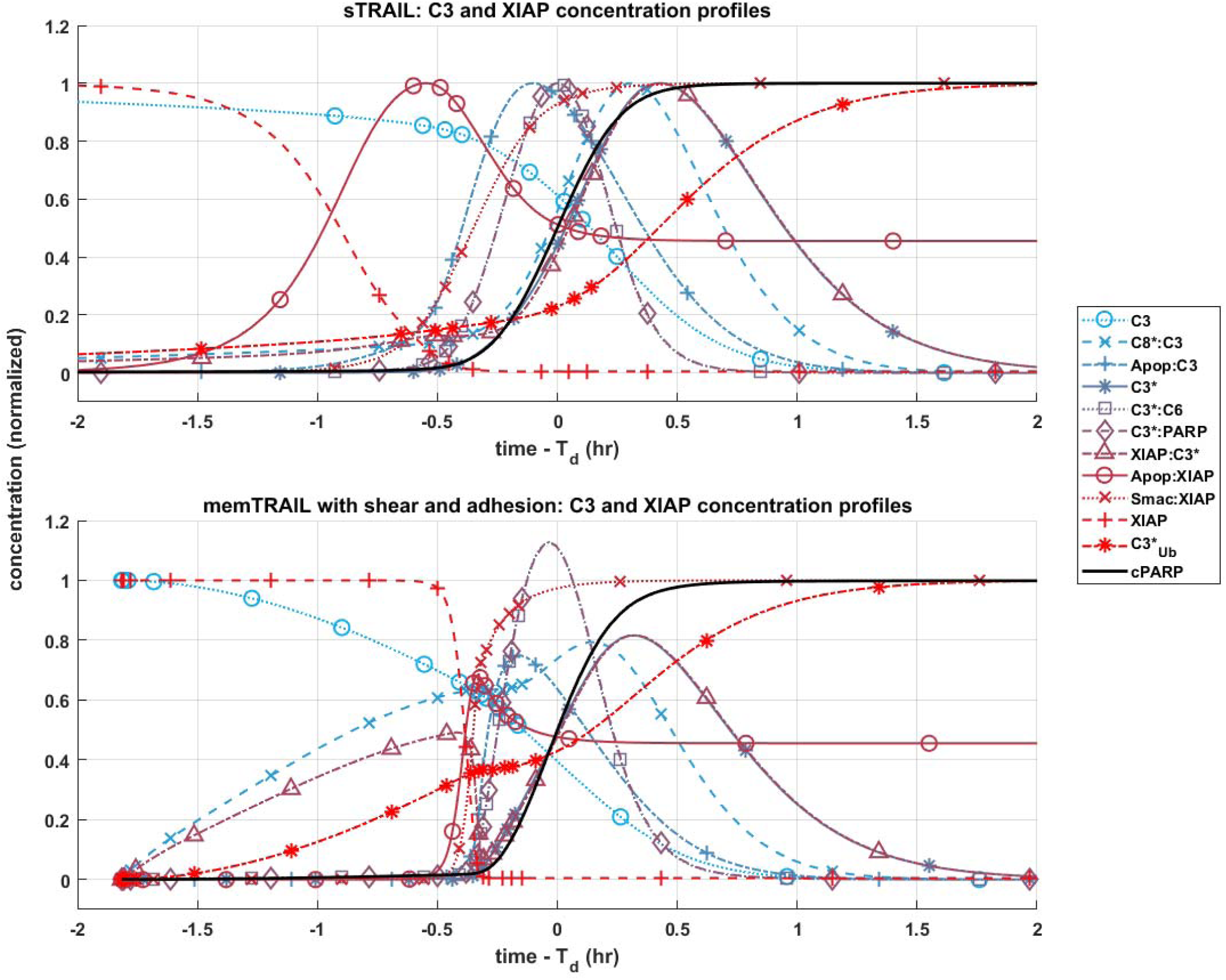
Post -mitochondrial pathway species concentration profiles as a function of time for four different cases of TRAIL bindng to DRs: sTRAIL, memTRAIL without shear, memTRAIL with shear without adhesion, and memTRAIL with shear and adhesion.

### 3.2 Mapping time until apoptosis

A sensitivity study was conducted to determine why liposome bound TRAIL (*memTRAIL*) acts as a more potent drug delivery mechanism. In this study, apoptosis was quantified by the time it takes for the variable *PARP* to cleave and form *cPARP*, which indicates the end of the reaction pathway from *TRAIL* binding to *DR* to eventual cell death. This point was defined as the time when *cPARP* was halfway through its transition time, 0.5 × *cPARP_max_*, *T_d_.* In order to gain insights about general trends, mappings of *T_d_* were created as a function of different combinations of the following parameters: Death Receptor (*DR*) concentration, *TRAIL* concentration, forward binding association rate constant of *TRAIL* binding *DR*, *k*_+_, and backwards binding dissociation rate constant, *k*_. Reasonable values of each parameter were determined, as specified in the Methods section, and then varied on a *log*_2_ scale.

#### 3.2.1 Varying: *TRAIL* and *DR* concentration

The first analysis considered a range of *DR* and *TRAIL* concentrations (Figure 5a). As *DR* concentration was increased, it was noted that *T_d_* decreased as expected. Interestingly, a bimodal dependence was observed, with *T_d_* starting high for lower concentrations of *TRAIL*, reaching a minimum at some intermediate value, and then trending upwards again at very high concentrations of *TRAIL* and low concentrations of *DR.*

**Figure 5.**
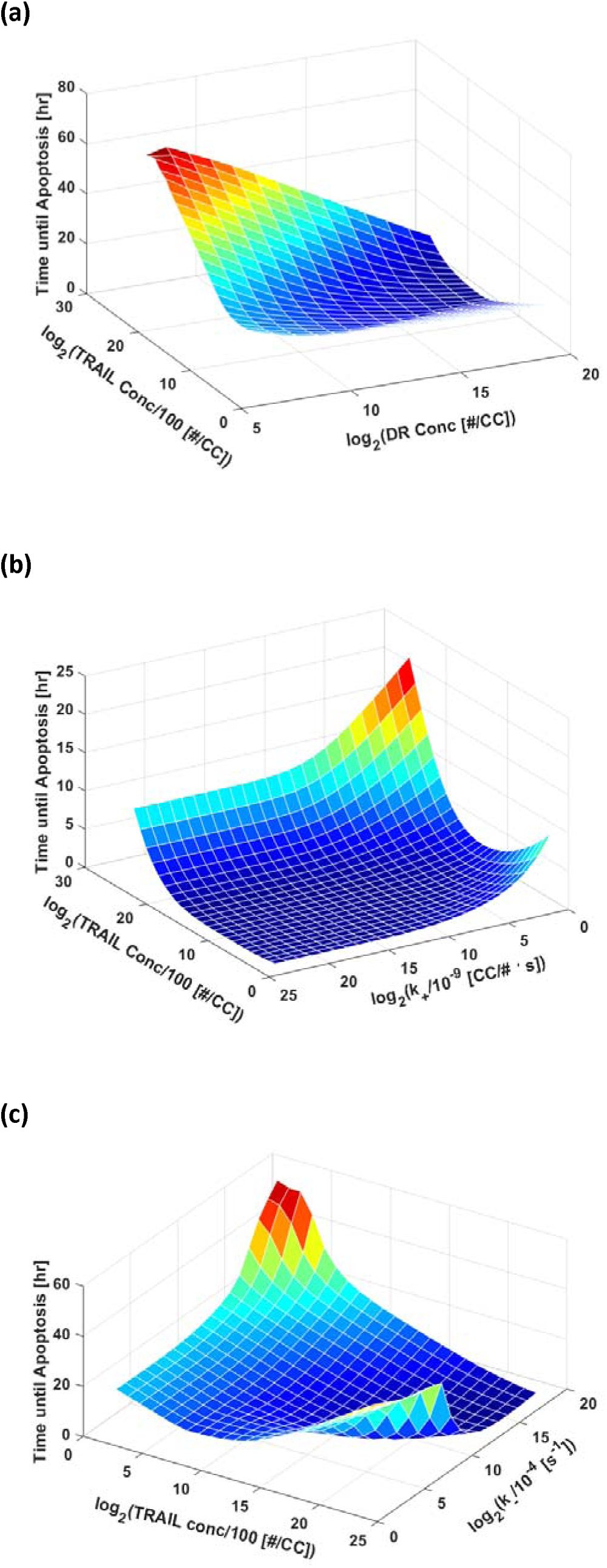
C3 and XIAP species concentration profiles as function of time for sTRAIL and memTRAIL with shear and adhesion.

#### 3.2.2 Varying: *TRAIL* concentration and *k*_+_

*TRAIL* concentration and *k_+_* were also varied (Figure 5b). As *k*_+_ was increased, time until apoptosis decreased, and a local minimum was found as a function of *TRAIL* concentration at low values of *k*_+_.

#### 3.2.3 Varying: *TRAIL* concentration and *k*_

*T_d_* was calculated as a function of *TRAIL* concentration and *k*_ (Figure 5c). As *k*_ was increased, *T_d_* increased; at very high values of *k*_, a characteristic decrease,and then increase in *T_d_* as a function of increasing *TRAIL* concentration was observed.

Taken together, these findings suggest that there is an optimum concentration of *TRAIL* that minimizes the time necessary for a cell to experience apoptosis.

## 4 Discussion

Previous experimental work has shown that leukocyte-tethered *TRAIL* (*memTRAlL*) is much more effective than soluble *TRAIL* (*sTRAIL*) at inducing apoptosis in CTCs (Mitchell et al., 2014). The model presented here offers an explanation. Given that both leukocytes and CTCs travel along similar streamlines in the blood flow and the high ratio of ∼1 × 10^6^ leukocytes per CTC, each CTC is expected to come into frequent contact with leukocytes throughout the vascular network (Yu et al., 2011). With this higher effective concentration of *memTRAIL* present in the vicinity of the CTCs, an elevated on-rate caused by shearing, and a reduced off-rate caused by *E* – *selectin* (*ES*) induced adhesion it was shown that *memTRAIL* induces apoptosis in under 2 *hr* while *sTRAIL* takes far longer at 10 + *hr .* These findings are consistent with those of Mitchell et al., and support the observation that treatment is much more effective when tethering the *ES/TRAIL* liposomes to leukocytes rather than relying on *sTRAIL* and its rapid clearance rate in the circulation (12, 14).

Given these findings, a deeper understanding was sought as to why the complete reaction proceeded more quickly, by varying the dynamics of *TRAIL* binding to *DRs.* First, four key parameters, *DR* concentration, *TRAIL* concentration, *k*_+_, and *k*_ were varied to determine the effect on the time until apoptosis, *T_d_*, after initial binding of *TRAIL* to *DR.* These simulations showed that given certain conditions, higher concentrations of *TRAIL* slow the overall reaction pathway. This suggests that the kinetics of *TRAIL* binding to *DRs* does not completely regulate how rapidly the overall apoptosis reaction proceeds, but rather affects how downstream reagents proceed in initiating cell death.

Four specific cases of binding were considered in greater detail: *sTRAIL*, *memTRAIL* without shear, *memTRAIL* with shear but without adhesion, and finally *memTRAIL* with shear and adhesion. For each of these cases, we looked at how key reagents unfold with respect to one another. For *memTRAIL*, pre-mitochondrial reagents started to transition immediately and completed their transition before *cPARP* transitioned. On the other hand, in considering the *sTRAIL* pathway to cell death, most of the pre-mitochondrial pathway reagents transitioned simultaneously with *cPARP*, which indicates that the pre-mitochondrial pathway is the limiting pathway for the *sTRAIL* case, but not for the *memTRAIL* with shear and adhesion case.

The relative concentrations of several key species between the two pathways were considered to focus in on the mechanism causing rapid apoptosis in CTCs treated in shear with *ES/TRAIL* liposomes. Due to the new configuration of *memTRAIL* conjugated to the surface of the liposome and the higher binding rate constants induced by shearing, the initial *memTRAIL* binding *DR* reaction occurred much faster, promoting the availability of its downstream reactants. The surge in concentration of one of these reactants, *C*8^∗^, pushed the reaction of *C*8^∗^ binding *C*3 forward rapidly, thus activating *C*3 to form *C*3^∗^.*XIAP* quickly engaged the available *C*3^∗^, as shown by the rapid increase in *XIAP: C*3^∗^ concentration but lack of increase in *C*3^∗^ alone. This increase leads to a critical difference between the two pathways. As *XIAP: C*3^∗^ concentration elevates, *C*3^∗^ is converted to
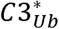
in an irreversible reaction, which decreases the amount of usable *C*3^∗^ until a critical point where *XIAP*: *C*3^∗^ is driven in the reverse direction to favor the unbound *C*3^∗^ and *XIAP* complex. This is marked by the sudden but temporary decrease in *XIAP*: *C*3^∗^ complex concentration. This new equilibrium point liberates more *C*3^∗^ for participating in the *memTRAIL* pathway than for the *sTRAIL* pathway, which promotes a steep and rapid uptake by *PARP* and *C*6 to form *C*3^∗^: *PARP* and *C*3^∗^: *C*6 respectively. Since *C*3^∗^:*PARP* peaks at a higher value, *PARP* is cleaved faster and apoptosis occurs more rapidly.

This sequence of events could also explain why extremely high values of *TRAIL* lead to higher *T_d_.* If the initial binding pathway proceeded too quickly, then
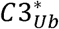
would consume too much of *C*3^∗^ from the reaction pathway before *C*3^∗^ was able to bind to *PARP*, retarding the reaction of *C*3^∗^ cleavage of *PARP.*

Thus, the efficiency of *ES/TRAIL* liposomes can be attributed to two effects. First, as the initial *TRAIL* binding *DR* pathway occurs more rapidly, the downstream reactions can begin sooner, leading to earlier apoptosis. The second effect leading to ES/TRAIL liposome efficiency is the intricate balance of
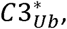
resetting the bound and unbound equilibrium of *C*3^∗^ and *XIAP.* While it is initially faster to have *C*3^∗^ and *XIAP* bind quickly to allow downstream reactions to begin, this increased rate can reach a point of diminishing returns, where too much *C*3^∗^ is irreversibly converted to
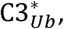
reducing the available *C*3^∗^ for downstream reactions which ultimately result in the cleavage of *PARP.*

## 5 Conclusion

These results faithfully recapitulate the experimental findings of Mitchell et al. (14)., where *sTRAIL* and tethered *ES/TRAIL* liposomes were both tested in mice to neutralize intravenously injected CTCs Given renal clearance mechanisms, our model suggests that although *sTRAIL* will eventually become effective if exposure could be sustained sufficiently long, it is unlikely to have sufficient time to act on the CTCs in vivo to induce apoptosis before being cleared from the circulation. This was the case in the previous experimental study of Mitchell et al. (CITE), as *sTRAIL* was ineffective at treating the mice whereas *ES/TRAIL* functionalized leukocytes showed 98.5% efficiency at clearing injected cancer cells after just 2 h of circulation. Given a new effective concentration of *memTRAIL*, an elevated *k*_+_ of *memTRAIL* binding *DR* caused by the slip velocity between the cell surfaces in shear flow, and a lowered off rate caused by *ES* adhesion, our simulation reproduces the behavior observed experimentally by Mitchell et. al, (14)and shed light on the enhanced efficacy of TRAIL/ES liposome therapy.

## 6 Methods

### 6.1 2D Binding: Initial Conditions

#### 6.1.1 Bound TRAIL

The model of Albeck et al. was modified, along with the provided initial conditions and rate constants, to capture the sheared liposomal TRAIL treatment of CTCs (16). First, it was necessary to determine the concentration of TRAIL bound to the surface of a liposome, to provide appropriate initial conditions for the model (16). Mitchell et al. estimated that there were ^∼^65 TRAIL molecules on average bound to the surface of each liposome of diameter ^∼^100 nm (14). For spherical liposomes, this corresponds to a surface density of liposomal TRAIL of *σ_L_* = 2 × 10^11^ *molecules/cm*^2^. The 2D density of death receptor on the surface of CTC was estimated, by assuming that all death receptors were located on the surface of the CTC. Given that there are 1 × 10^4^ *DRA/cellular volume* and the volume of the cell to is 1 × 10^−9^*cm*^3^, for a roughly spherical cell the surface density can be calculated to be *σ_R_* = 2 × 10^−9^*molecules/cm*^2^.

#### 6.1.2 Soluble TRAIL

An effective spatial availability of *sTRAIL* was determined that corresponds to conditions in the previous experiments of Mitchell et al. (14). Plasma concentration of *sTRAIL* was estimated to be 1 *μg/mL.* For a cellular volume of 1 × l0^−9^*cm*^2^, it was approximated that the concentration of *sTRAIL* available for surface reaction is is 1.85 × 10^4^ *TRAIL/ cellular volume*, or more appropriately, 3.8 × 10^−9^ *TRAIL/cm*^2^.

### 6.2 2D Binding: Reaction Rate Constants

#### 6.2.1 Binding Association Rate Constant

##### 6.2.1.1 Chang and Hammer Approach: Shear

Next, the original 3D model was modified to more appropriately represent the binding kinetics of liposomal TRAIL in shear, via the binding association and dissociation rate constants.

An analysis was carried out employing Chang and Hammer’s approach for determining binding association and dissociation rates for a 2D surface binding to a 2D surface with a relative slip velocity between the surfaces (25). The first step in this analysis required a determination of the slip velocity between liposomal TRAIL attached to leukocytes and CTCs in shear. It was assumed that the centroid of a TRAIL functionalized leukocyte and the centroid of an interacting CTC were ^∼^10 *μm* apart (the sum of their radii), *d*, when they convect past each other. Given a uniform shear rate, *S*, of 1000 as in typical blood flow (26), it was determined that the relative velocity of the centers was *S* × *d* = ***V*** = 1 *cm/s.* This slip velocity, the sum of the lateral diffusivities of TRAIL and death receptor, and the reactive radius of our reagents were used to determine the Peclet number, *Pe* = ***V*** · *a/D*, which is the dimensionless ratio of bulk flow (advection) to diffusive flow of reagent. A diffusivity, *D*, was used of order 1 × 10^−9^*cm*^2^*/s* (27) and a reactive radius, *a*, of 5 × 10^6^*cm* (28) to yield a *Pe* of 5000. In Chang and Hammer’s analysis, since *Pe* ≫ 1, the Nusselt number, another measure of bulk flow to diffusive flow, can be approximated as *Nu* = 2*Pe/π*, yielding a *Nu* of 3183. From this, the enhanced forward association rate constant was determined to be *k_o_* = *πDNu* = 1.0 × 10^−5^ *cm*^2^*/s.*

##### 6.2.1.2 Chang and Hammer Approach: Unsheared (Diffusion Limit)

It also was necessary to determine the forward association rate constant, again in 2D binding, for the unsheared case to apply the model to the static control conditions in the experiments of Mitchell et al. This simulation condition involved no slip velocity, and thus *Pe* = 0. When *Pe* = 0, *Nu* = 2/log (^*b*^/_*a*_), where *b*1/2 the mean distance between ligand and receptor and *a* is the reactive radius, *b* was estimated to be of order of magnitude *b* = 10 × 10^−6^ *cm*, yielding *Nu* = 2.9 and *k_o_* = 9.1 × 10^−9^ *cm*^2^*/s*.

##### 6.2.1.3 Bell Approach: Unsheared (Diffusion Limit)

As a check on the assumptions made, Bell’s approach, which does not take into account a relative slip velocity, was also considered (29). Bell’s approach proposes that *k_o_* = 2*πD*, giving us *k_o_* = 6.3 × 10^−9^*cm*^2^*/s*. This value is of the same order of magnitude for the unsheared case using Chang and Hammer’s approach.

##### 6.2.1.4 Modification of Albeck Approach

Albeck, and others have measured a forward association rate constant of *sTRAIL* binding death receptor with value equal to 2.4 × 10^5^*M*^−1^*s*^−1^. To be consistent with the dimensionality of the molecular participants described above in 2D Binding: Initial Conditions, this value was converted to 1.94 × 10^−12^*cm*^2^/*s*, which directly follows from a CTC volume of 1 × 10^−9^*cm*^2^ and surface area of 4.8 × 10^−6^*cm*^2^.

#### 6.2.2 Binding Dissociation Rate Constant

##### 6.2.2.1 Chang and Hammer Approach: Sheared

To determine the off rate for the sheared case, the average duration of encounter, *τ*, which for *Pe* ≫ 1, can be approximated as *≫* ∼8*a*/(3|***V***|*π*), was estimated. Again, *a* is the reactive radius and |***V***| is the slip velocity, yielding *τ* ∼ 4.2 × 10^−6^*S*. Next, the dimensionless duration time, Λ = *τ*/(^*a*^2^^/_*D*_)) = 1.7 × 10^−4^, and the dimensionless Damköhler number, *δ* = *a*^2^*k_in_/D* = 2.5 × 10^7^, were determined, where *k_in_* is the intrinsic forward reaction rate, equal to 1 × 10^9^*s*^−1^ (28). Given these two parameters, the probability of binding was expressed as *P* = Λ*δ*/(1 + Λ*δ*) ∼ .9998. From this, the overall forward rate of reaction was found to be *k_f_* = *k_o_P* = 9.998 × 10^−6^ *cm*^2^/*s.* Given that *k_f_* = *k_o_k_in_*/(*k_in_* + *k*_), *k*_ was found to be *k*_ = 2.4 × 10^5^*s*^−1^.

##### 6.2.2.2 Chang and Hammer Approach: Unsheared (Diffusion Limit)

The binding dissociation rate constant was estimated for the unsheared case. Here, *τ* is given as *a*^2^/8*D*, yielding *τ* = 3.1 × 10^−3^ *s*, Λ = 0.125, *δ* = 2.5 × 10^7^, *P* ∼ 1 for a *k_f_* = 9.1 × 10∼^−9^*cm*^2^/*s* and a *k*_ = 320 *s*^−1^.

##### 6.2.2.3 Bell Approach: Unsheared (Diffusion Limit)

To check the assumptions made, Bell’s approach was referenced, which does not take into account a relative slip velocity. Bells approach states that *k*_ = 2*D/a*^2^, yielding *k*_ = 80 *s*^−1^. This value is within one order of magnitude for the unsheared case obtained separately via Chang and Hammer’s approach.

#### 6.2.3 E-selectin Effects on Binding Adhesion

Mitchell et al. demonstrated E-selectin as an adhesive targeting protein that simultaneously promotes the establishment of TRAIL-functionalized leukocytes, as well as close surface interactions with CTCs. To represent this adhesive interactions between TRAIL-coated leukocytes and colliding CTCs, the duration of time that E-selectin was expected to be adhered to CTCs before separating in shear flow was estimated to be of the order 1 × 10^−3^*S* (30). During this time, E-selectin tethering was assumed to effectively reduce the slip velocity between the two cells to zero, and thus set the binding dissociation rate to the diffusion limited case (unsheared). This was implemented within the simulation by executing the numerical integration for 1 × 10^−3^*s* with *σ_L_* as the initial condition. Following this interval, the numerical integration using the last concentration from the first interval as the initial condition for all reagents except *TRAIL* which was then set to zero.

## 8 Tables

### Summary of parameters

Table 1 lists the parameter values used in the different simulation conditions.

**Table 1.** Table of values used in simulation of TRAIL binding to DRs Sources: Albeck (16), Mitchell (14), Chang (25), Szegezdi (31)

|  | **TRAIL**<br>**Concentration,**<br>**$\sigma_L$** | **Death**<br>**Receptor**<br>**Concentration,**<br>**$\sigma_R$** | **Binding**<br>**Association**<br>**Rate Constant,**<br>**$k_o$** | **Binding**<br>**Dissociation**<br>**Rate Constant**<br>**$k_-$** | **Duration of**<br>**TRAIL**<br>**Exposure** |
| --- | --- | --- | --- | --- | --- |
| **Soluble TRAIL** | Albeck<br>(modified)<br>$3.8 \times 10^9$<br>[$\#/cm^2$] | Szegezdi<br>$2 \times 10^4$<br>[$\#/cm^2$] | Albeck<br>(modified)<br>$1.94 \times 10^{-12}$<br>[$cm^2/(\# \cdot s)$] | Albeck<br>$1 \times 10^{-3}$<br>[$s^{-1}$] | Continuous |
| **Liposomal**<br>**TRAIL, no**<br>**shear, with E-**<br>**selectin** | Mitchell<br>(modified)<br>$2 \times 10^{11}$<br>[$\#/cm^2$] | Szegezdi<br>(modified)<br>$2 \times 10^9$<br>[$\#/cm^2$] | Chang and<br>Hammer (no<br>slip velocity)<br>$9.1 \times 10^{-9}$<br>[$cm^2/(\# \cdot s)$] | Chang and<br>Hammer (no<br>slip velocity)<br>320<br>[$s^{-1}$] | Continuous |
| **Liposomal**<br>**TRAIL, shear,**<br>**with E-selectin**<br>**(assuming cell**<br>**adhesion)** | Mitchell<br>(modified)<br>$2 \times 10^{11}$<br>[$\#/cm^2$] | Szegezdi<br>(modified)<br>$2 \times 10^9$<br>[$\#/cm^2$] | Chang and<br>Hammer (slip<br>velocity)<br>$1 \times 10^{-5}$<br>[$cm^2/(\# \cdot s)$] | Chang and<br>Hammer (no<br>slip velocity)<br>320<br>[$s^{-1}$] | $1 \times 10^{-3}$ [s] |
| **Liposomal**<br>**TRAIL,**<br>**shear,**<br>**without E-**<br>**selectin** | Mitchell<br>(modified)<br>$2 \times 10^{11}$<br>[$\#/cm^2$] | Szegezdi<br>(modified)<br>$2 \times 10^9$<br>[$\#/cm^2$] | Chang and<br>Hammer (slip<br>velocity)<br>$1 \times 10^{-5}$<br>[$cm^2/(\# \cdot s)$] | Chang and<br>Hammer<br>$2.4 \times 10^5$<br>[$s^{-1}$] | Continuous |

## 10 Declarations

### 10.1 Ethics approval and consent to participate

Not Applicable

### 10.2 Consent to publish

Not Applicable

### 10.3 Availability of data and materials

Code for simulation available

### 10.4 Competing Interests

Not applicable

### 10.5 Funding

This work was funded by NIH Grant CA203991 to M.R.K.

### 10.6 Authors’Contributions

EL designed, implemented and evaluated the algorithmic methods and suggested improvements of the model. MK conceived the model and provided input to the algorithms’ development. EL wrote the manuscript. All authors edited the manuscript, read and approved the final manuscript.

## 10.7 Acknowledgements

Not applicable

